# *PCSK9* genetic variants, life-long lowering of LDL-cholesterol and cognition: a large-scale Mendelian randomization study

**DOI:** 10.1101/335877

**Authors:** Donald M. Lyall, Joey Ward, Maciej Banach, George Davey Smith, Jason G. Gill, Jill P. Pell, Michael V Holmes, Naveed Sattar

## Abstract

**Aims:** PCSK9 inhibitors lower LDL cholesterol and are efficacious at reducing risk of vascular disease, however questions remain about potential adverse effects on cognitive function. We examined the association of LDL cholesterol-lowering genetic variants in *PCSK9* with continuous measures of cognitive ability

**Methods and Results:** Six independent SNPs in *PCSK9* were used in up to 337,348 individuals from the UK Biobank who underwent measures of cognitive ability (fluid reasoning, reaction time, trial making test and digit symbol coding. Scaled to a 50mg/dL lower LDL cholesterol, the *PCSK9* allele score was associated with a lower risk of CHD (odds ratio 0.73; 95% CI: 0.60 to 0.90, P = 0.003). The scaled *PCSK9* allele score nominally associated with worse log reaction time (0.04 standard deviations; 95%CI: 0.00, 0.08; P=0.038). Although no strong associations of the *PCSK9* allele score were identified with any cognitive trait, the imprecision around the estimates meant that we could not exclude a similar magnitude of effect of genetic inhibition of PCSK9 to that seen with established risk factors, including *APOE*e4 or smoking status for any of the individual cognition traits. Point estimates for the *PCSK9* allele score and cognition traits were all on the harmful side of unity.

**Conclusions:** Using currently available data in UK Biobank, we are not able to rule out meaningful associations of *PCSK9* genetic variants with cognition traits. These data highlight the need for further large-scale genetic analyses and, in parallel, continued pharmacovigilance for patients currently treated with PCSK9 inhibitors.

## Introduction

Therapeutic modification of atherogenic lipoproteins by statins^1^, ezetimibe^2^ and Proprotein convertase subtilisin/kexin type 9 (PCSK9) inhibition^3^ is an effective strategy to reduce risk of cardiovascular disease (CVD), principally by lowering non-HDL-cholesterol^4^. PCSK9 regulates LDL-cholesterol (LDL-C) through hepatic expression of LDL receptors. PCSK9 inhibitors are licensed as LDL-C lowering agents with excellent efficacy and evidence of cardiovascular benefit: the FOURIER trial^3^ recently reported that a monoclonal antibody to PCSK9 that lowered LDL-C by 59% (∼56 mg/dl) led to a 15% reduction in risk of a composite of cardiovascular death, myocardial infarction, stroke, hospitalization for unstable angina or coronary revascularization in patients with established atherosclerotic vascular disease. However, early phase 3 studies showed a potential excess of neurocognitive adverse effects ^5^, leading the US FDA to instruct pharmaceutical companies to assess neurocognitive side effects of PCSK9 inhibitors. While, reassuringly, no excess risk was noted in the FOURIER trial^3^ or its substudy with detailed cognitive measures^6^, these studies cannot exclude potential adverse effects from longer-term use of PCSK9 inhibitors. An orthogonal approach to obtain reliable information is to exploit genetic variants that mimic pharmacological inhibition of PCSK9. Naturally occurring variation in the gene encoding a drug target can be used to gauge insight on long-term effects of therapeutic modification. So-called drug-target Mendelian randomization^7^ exploits the characteristics of genotype for the reliable estimation of both intended and unintended consequences of therapeutic modification of a drug target, as previously demonstrated^8-11^.

## Methods

We used four measures of cognitive ability from UK Biobank (UKB) data: two from baseline (2006-2010): fluid reasoning measured in 160,130 individuals (with genetic data, prior to exclusions listed below) and reaction time in 482,187 as they showed good intra-participant longitudinal reliability in n=19,999 participants^12^, and two measured between 2014-15 via the internet: trail making test (TMT) A (processing speed) in 100,587 and B (speed plus executive function) in 100,610, and digit symbol coding (executive function) in 115,933. TMT and reaction time scores were log-transformed due to a positive skew. All measures were standardized to Z-scores.

Six independent SNPs (R^2^<0.15) in *PCSK9* (referent allele frequencies rs2479394 A = 0.72; rs11206510 C = 0.19; rs2479409 A = 0.65; rs10888897 T = 0.39; rs7552841 C = 0.63; rs562556 G = 0.18); orientated so that the effect allele associated with a lower LDL-C were used as genetic variants to proxy therapeutic inhibition of PCSK9. SNPs were selected from the paper by Ference and colleagues in NEJM^13^. In the paper by Ference^13^, seven *PCSK9* SNPs were used, however two were found to be moderately correlated (rs2479409 and rs2149041; R^2^=0.37). We therefore removed the SNP with the weaker association with LDL-C (rs2149041) as defined by the P-value^13^, leaving six SNPs (with pair-wise LD R^2^<0.15) described above. The LDL-C association of each of the six *PCSK9* SNPs from the Global Lipids Genetics Consortium^14^ was used to construct a weighted *PCSK9* allele score. Linear regression analyses used the four cognition traits as dependent variables, the weighted *PCSK9* score as the independent variable, adjusted for age, sex, GWAS array, and 10 principal components (as provided by UKB). To mimic pharmacological modification of *PCSK9*, we report results of the *PCSK9* allele score scaled to the 50mg/dL lower LDL-C achieved in FOURIER. We compared estimates of the *PCSK9* allele score with cognition traits to those from *APOE* e4 dosage (excluding rare *APOE* e2/e4), *APOE* e4/e4 vs. e3/e3 homozygosity; current vs. never smoking, and 5-years of increased cross-sectional age with cognitive traits. As a further positive control, we tested the association of the PCSK9 allele score with risk of CHD in UK Biobank (defined as self-reported physician-diagnosed myocardial infarction and angina).

We excluded participants with non-white British ancestry, self-report vs. genetic sex mismatch, putative sex chromosomal aneuploidy, excess heterozygosity, and missingness rate >0.1. We removed one random participant in cases where two individuals were first cousins or closer. Stata v14 and PLINK v1.90 were used for analyses.

## Results

The *PCSK9* allele score scaled to a 50mg/dL lower LDL-C was associated with lower risk of CHD in UKB (comprising 15,284 cases of myocardial infarction and angina in 338,852 individuals; odds ratio: 0.73; 95% CI: 0.60 to 0.90, P = 0.003).

We next investigated the associations of six variants in *PCSK9* scaled to 50mg/dL lower LDL-C in 109,870 individuals with measures of fluid reasoning, 337,348 with reaction time, 73,044 with processing speed (TMT A), 73,063 with processing speed plus executive function (TMT B), and 82,012 with digit symbol coding (executive function).

The *PCSK9* allele score was nominally associated with log reaction time (0.04 SDs higher log reaction time; 95%CI: 0.002, 0.079; P=0.038). For fluid reasoning, the scaled *PCSK9* allele score had wide 95% CI (−0.08, 0.07) that included the estimates for the association of 5 years additional age (−0.05 SDs, 95%CI: −0.06, −0.05). Similar patterns were identified for all other cognition traits, meaning that despite the large sample size, the imprecision around the estimates obtained from the *PCSK9* allele score meant that we could not exclude a similar magnitude of effect of genetic inhibition of PCSK9 to that seen with the positive controls, including *APOE*e4 or smoking status for any of the individual cognition traits (**Figure**). Notably, point estimates for the association of the *PCSK9* allele score and all cognitive ability end-points were on the harmful side of unity.

**Figure.**
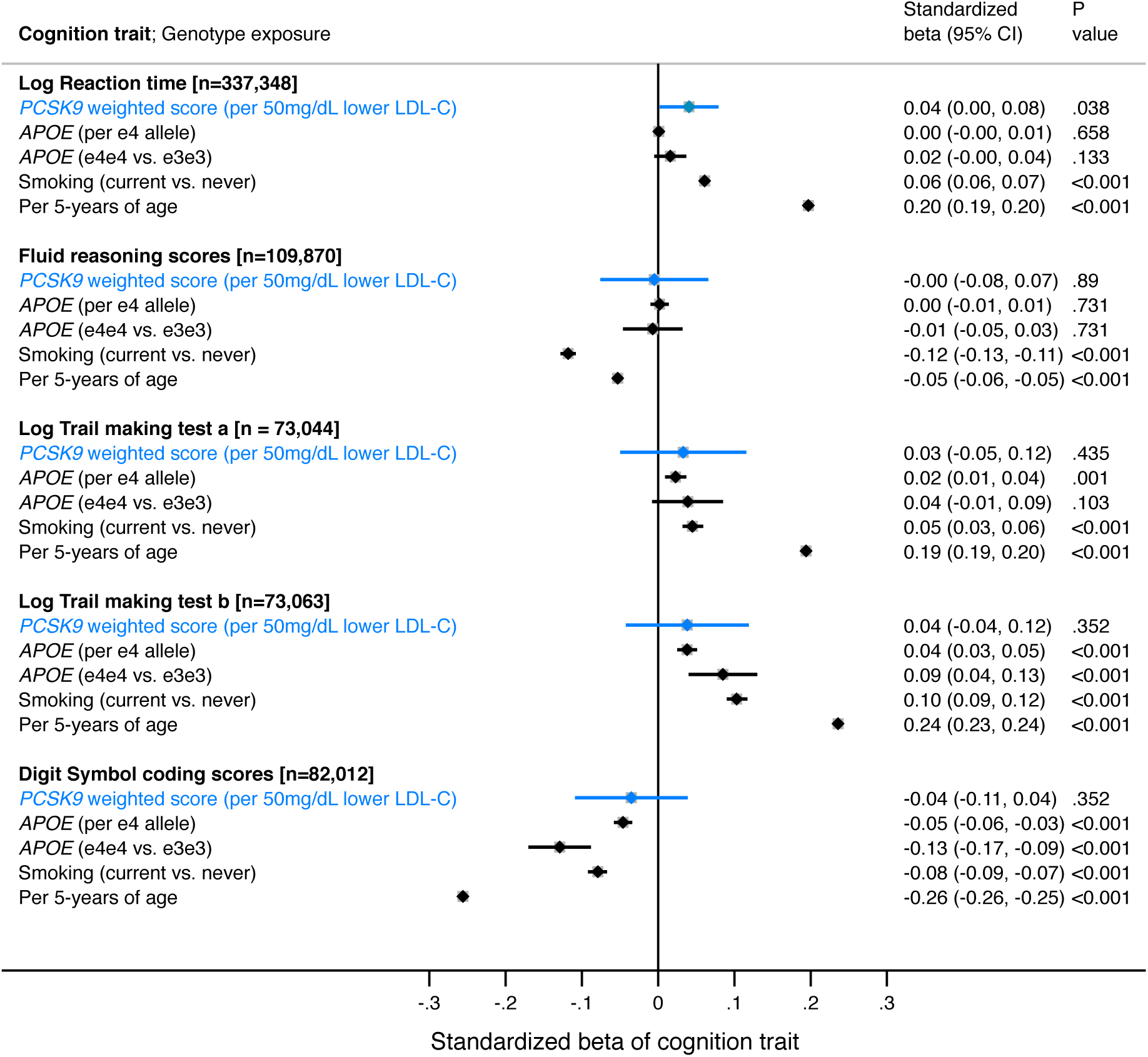
Association of a *PCSK9* allele score (in blue) scaled to 50mg/dL lower LDL-cholesterol and other selected exposures (in black) with cognitive traits in UK Biobank.

In sensitivity analyses, removal of participants that self-reported a neurological condition (∼5% of the dataset^ref12^) did not alter the findings, nor did substituting rs2479409 for rs2149041 in the *PCSK9* allele score.

## Discussion

In this large-scale analysis of individuals from the general population, we used naturally occurring genetic variants in *PCSK9* to gauge insight into the effect of lifelong lowering of LDL-C through inhibition of PCSK9 and its association with cognition abilities. Using currently available data in UKB, we were not able to provide definitive evidence on the relationship of *PCSK9* genetic variants with cognition traits. While this may have arisen due to lack of power, we note that we were able to show associations of the PCSK9 allele score with risk of prevalent self-reported CHD, and also we note robust associations of conventional risk factors (e.g. age and smoking) and genetic variants (e.g. *APOE* e4) with the cognitive traits. While the imprecision makes it challenging to draw firm conclusions about an effect (or lack thereof) of life-long LDL-cholesterol lowering by PCSK9 inhibition on cognition, our data highlight the need for additional large-scale genetic analyses. In parallel, continued pharmacovigilance is needed for patients currently treated with PCSK9 inhibitors.

## Acknowledgements

This research was conducted using UK Biobank application 17689. MVH works in a unit that receives funds from the University of Oxford and the UK Medical Research Council and is supported by a British Heart Foundation Intermediate Clinical Research Fellowship (FS/18/23/33512) and the National Institute for Health Research Oxford Biomedical Research Centre. GDS works in a unit that receives funds from the University of Bristol and the UK Medical Research Council (MC_UU_12013/1). The funders had no role in study design, decision to publish, or preparation of the manuscript. UK Biobank was established by the Wellcome Trust medical charity, Medical Research Council (MRC), Department of Health, Scottish Government and the Northwest Regional Development Agency. We are grateful to UK Biobank participants.

## Disclosures

The Clinical Trial Service Unit & Epidemiological Studies Unit at the University of Oxford (MVH) has received research grants from Abbott/ Solvay/Mylan, AstraZeneca, Bayer, GlaxoSmithKline, Merck, Novartis, Pfizer, Roche, and Schering. MVH has collaborated with Boehringer Ingelheim in research, and in accordance with the policy of The Clinical Trial Service Unit and Epidemiological Studies Unit (University of Oxford), did not accept any personal payment. NS has consulted for AstraZeneca, Bristol-Myers Squibb, Amgen, Sanofi, and Boehringer Ingelheim. No other authors reported disclosures.

